# Archaeal extracellular vesicles are produced in an ESCRT-dependent manner and promote gene transfer and nutrient cycling in extreme environments

**DOI:** 10.1101/2021.02.09.430445

**Authors:** Junfeng Liu, Virginija Cvirkaite-Krupovic, Pierre-Henri Commere, Yunfeng Yang, Fan Zhou, Patrick Forterre, Yulong Shen, Mart Krupovic

## Abstract

Membrane-bound extracellular vesicles (EVs), secreted by cells from all three domains of life, transport various molecules and act as agents of intercellular communication in diverse environments. Here we demonstrate that EVs produced by a hyperthermophilic and acidophilic archaeon *Sulfolobus islandicus* carry not only diverse proteins but also chromosomal and plasmid DNA, and can transfer this DNA to recipient cells. Furthermore, we show that EVs can support the heterotrophic growth of *Sulfolobus* in minimal medium, implicating EVs in carbon and nitrogen fluxes in extreme environments. Finally, our results suggest that, similar to eukaryotes, production of EVs in *S. islandicus* depends on the archaeal ESCRT machinery. Using synchronized *S. islandicus* cultures, we show that EV production is linked to cell division and appears to be triggered by increased expression of ESCRT proteins during this cell cycle phase. Using a CRISPR-based knockdown system, we show that archaeal ESCRT-III and AAA+ ATPase Vps4 are required for EV production, whereas archaea-specific component CdvA appears to be dispensable. Collectively, our results suggest that ESCRT-mediated EV biogenesis has deep evolutionary roots, likely predating the divergence of eukaryotes and archaea, and that EVs play an important role in horizontal gene transfer and nutrient cycling in extreme environments.

## INTRODUCTION

Extracellular vesicles (EVs) are membrane-bound particles of variable diameter secreted by the cells into extracellular milieu. Although known for several decades, EVs were broadly regarded as cellular waste products, debris or artifacts of lipid aggregation [1]. However, the growing body of data shows that EVs play multiple, biologically important roles in all three domains of life [1-7]. During the past decade, it was discovered that EVs are responsible for intercellular shuttling of diverse cargoes, including proteins, DNA, RNA, lipids and various signaling molecules [5,7-10], and may promote certain human pathologies [11,12], including cancer [13,14]. Furthermore, EVs hold great promise as vehicles for drug targeting and delivery [15,16]. Finally, EVs may play an important ecological role, especially, in aquatic ecosystems [5]. It has been shown that DNA-carrying EVs produced by diverse bacteria, including *Prochlorococcus*, a numerically dominant marine cyanobacterium, are abundant in coastal and open-ocean seawater samples [7]. Importantly, *Prochlorococcus* EVs could support the growth of heterotrophic bacterial cultures, which implicates EVs in marine carbon flux [7]. Archaea of the phyla Euryarchaeota and Crenarchaeota are also known to produce EVs under laboratory conditions. *Thermococcus prieurii*, but not other *Thermococcus* species, secrete EVs packed with elemental sulfur, presumably to prevent the accumulation of toxic levels of sulfur in the cytoplasm [17]. Furthermore, similar to bacteria, EVs produced by members of the phylum Euryarchaeota thriving in deep-sea hydrothermal vents (order Thermococcales) and saline lakes (order Halobacteriales) were shown to carry DNA [18-21]. Whether the same is true for EVs produced by crenarchaea of the order Sulfolobales, which represent major inhabitants of terrestrial acidic hot springs, remains unknown. It is also unclear whether archaeal EVs are secreted under natural growth conditions in the environment.

The mechanisms of EV biogenesis have been extensively studied in eukaryotes but remain poorly understood in bacteria and archaea [1,3]. In eukaryotes, the most studied mechanism of EV formation relies on the endosomal sorting complex required for transport (ESCRT) machinery [22-24]. Many archaea also encode homologs of the ESCRT system but its involvement in EV biogenesis remains unclear. The ESCRT machinery is responsible for many key membrane remodeling processes in eukaryotic cells, including membrane abscission during cytokinesis, biogenesis of certain types of EVs and multivesicular bodies, and budding of enveloped viruses, such as HIV-1 and Ebola virus [22,25,26]. The ESCRT proteins assemble on the cytosolic face of the membrane and drive membrane bending and scission reaction [26]. The ESCRT machinery can be subdivided into several functionally distinct subcomplexes known as ESCRT-0, ESCRT-I, ESCRT-II and ESCRT-III as well as AAA+ ATPase Vps4. Among these, ESCRT-III and Vps4 are universally involved in ESCRT-dependent membrane remodeling processes, whereas ESCRT-0, ESCRT-I and ESCRT-II are compartment-specific and facilitate recruitment of ESCRT-III to diverse membranes in different cellular contexts [25,27]. ESCRT-III proteins form a ring-like filament at the membrane, whereas the Vps4 ATPase binds directly to ESCRT-III and dynamically disassembles the ESCRT-III complex in an ATP-dependent manner, thereby driving membrane-remodeling [28,29].

Similar to eukaryotes, most archaea of the TACK (for Thaumarchaeota, Aigarchaeota, Crenarchaeota and Korarchaeota) and Asgard superphyla encode an ESCRT machinery [30-33]. Interestingly, the ESCRT machinery encoded by Asgard archaea is phylogenetically more closely related to the eukaryotic homologs compared to those from other archaea and Asgard Vps4 could efficiently complement the *vps4* null mutant of *Saccharomyces cerevisiae* [31,34]. However, due to difficulties in cultivation and lack of genetic tools in Asgard archaea, the role of their ESCRT machinery in membrane remodeling remains to be investigated in this superphylum. The archaeal ESCRT system has been experimentally investigated in *Sulfolobus* and *Nitrosopumilus* species [33,35-41], with *Sulfolobus* representing the model organism for elucidation of the role and functioning of the archaeal ESCRT machinery [30,41,42]. In hyperthermophilic archaea of the order Sulfolobales, the ESCRT machinery is the key component of cell division apparatus composed of AAA+ ATPase Vps4 (also known as CdvC), four ESCRT-III homologs (ESCRT-III [CdvB], ESCRT-III-1 [CdvB1], ESCRT-III-2 [CdvB2], ESCRT-III-3 [CdvB3]), and archaea-specific component CdvA. The latter protein binds to DNA [43,44] and recruits ESCRT-III to the membrane [44]. CdvA is not homologous to the eukaryotic ESCRT-0, ESCRT-I or ESCRT-II [45], and is missing in certain archaea, including some thaumarchaea [46] and aigarchaea [47]. It has been also proposed that the *Sulfolobus* ESCRT machinery is involved in viral assembly within the cytoplasm and in escape from the infected cell by using a unique lysis mechanism [48]. Whether the function of archaeal ESCRT machinery can be extended to other membrane remodeling processes remains to be demonstrated. Notably, ESCRT-III-1, ESCRT-III-2 and Vps4 were identified among proteins present within EVs secreted by *Sulfolobus acidocaldarius, S. solfataricus* and *S. tokodaii* [49]. This finding suggested that ESCRT machinery is involved in EV biogenesis [49]. However, EVs are known to be produced by archaea which lack the functional ESCRT system and divide using the bacterial-like FtsZ-based cell division machinery, including halophilic archaea (class Halobacteria) and members of the order Thermococcales [18,50].

Here we characterize the composition, role and biogenesis of EVs produced by a hyperthermophilic and acidophilic archaeon *Sulfolobus islandicus*. We demonstrate that besides proteins, *Sulfolobus* EVs carry chromosomal and plasmid DNA, and that EVs can transfer this DNA to recipient cells. We also investigate the role of the *Sulfolobus* ESCRT machinery in EV biogenesis and show that all four ESCRT-III homologs and Vps4 ATPase play an important role in this process, whereas CdvA appears to be dispensable. Using synchronized *S. islandicus* cultures, we demonstrate that EV production is linked to cell division and appears to be triggered by the cell cycle-coordinated fluctuations in the expression of ESCRT proteins. Finally, we show that EVs similar to those produced by *Sulfolobus* cells under laboratory conditions can be also found in the environmental sample. Collectively, our results suggest that the ESCRT-dependent mechanism of EV biogenesis is conserved in the archaeo-eukaryotic lineage and that EVs play an important role in gene transfer in extreme environments.

## MATERIALS AND METHODS

### Strains, growth conditions and transformation of *Sulfolobus*

*Sulfolobus islandicus* strains REY15A and E233S (REY15A*ΔpyrEFΔlacS*) [51], and *Sulfolobus solfataricus* PH1-16 (PH1 *pyrF* mutant) [52], hereafter PH1-16, were grown aerobically with shaking (145 rpm) at 75°C in 30 ml of STVU medium containing mineral salts (M), 0.2% (wt/vol) sucrose (S), 0.2% (wt/vol) tryptone (T), a mixed vitamin solution (V) and 0.01% (wt/vol) uracil (U); the pH was adjusted to 3.5 with sulfuric acid, as described previously [51]. SCV medium containing 0.2% (wt/vol) casamino acids (C) was used for selection of uracil prototrophic transformants. ATV medium containing 0.2% (wt/vol) arabinose (A) was used to induce protein overexpression and RNA interference. The plasmids and strains constructed and used in this study are listed in Tables S2 and S3, respectively.

### Isolation and purification of EVs

EVs were isolated from liquid cultures of *S. islandicus* E233S or *S. solfataricus* PH1-16 strains carrying shuttle vector pSeSD. The cells were grown at 75°C in appropriate medium and EVs were harvested at the indicated times. Cells were removed by centrifugation at 7,000 rpm at 4°C for 20 min. The supernatant was filtered with 0.45 μm filter and EVs collected by ultracentrifugation at 40,000 rpm (Type 45 Ti rotor) at 4°C for 2 h, followed by 100,000 rpm (TLA 100.2 rotor) at 4°C for 1 h, and then re-suspended in 500 μl of PBS.

For mass spectrometry and DNA content sequencing, the EVs were collected during the exponential growth phase (24 h) and further purified by ultracentrifugation in sucrose gradient (50%, 45%, 40%, 35%, 30%, and 25%) at 25,000 rpm (SW 60 rotor) at 4°C for 10 min. The EVs formed an opalescent band in the region of the gradient corresponding to 30-40% sucrose (Fig. S15). The band was collected and EVs pelleted by ultracentrifugation at 100,000 rpm (TLA100.2 rotor) at 4°C for 1 h. The resulting pellet was re-suspended in 500 μl of PBS.

### Transmission electron microscopy and determination of the relative EV diameters

For TEM analysis, EVs or cell cultures were absorbed to glow-discharged copper grids with carbon-coated Formvar film and negatively stained with 2.0% (w/v) uranyl acetate. The samples were observed under FEI Tecnai Spirit BioTwin 120 microscope (FEI, Einthoven, The Netherlands) operated at 120 kV.

Due to pleomorphicity of EVs, their relative diameters were determined by measuring the corresponding area (A) using ImageJ. First, the electron micrographs of negatively stained EVs were opened in ImageJ and the scale of each image was set according to the scale bar in the corresponding micrograph. Then the area of each EV was measured separately and the relative diameter (D) was calculated according to the area, based on the equation A=(π/4) × D^2^.

### Detection of EVs in environmental samples

Hot spring water samples collected from the solfataric field of the Campi Flegrei volcano in Pozzuoli, Italy have been described previously [53]. Twelve milliliters of the sample were concentrated by ultracentrifugation at 38,000 rpm (SW41 rotor) at 15°C for 3 h. After centrifugation, most of the supernatant was removed and the pellet was resuspended in the residual liquid (∼300 μl). The concentrated sample was prepared for TEM analysis as described above.

### Flow cytometry and quantification of EVs

EVs were isolated from 50 ml cultures of E233S and PH1-16 cells carrying pSeSD vector at the given time points of cell growth, then 50 μl of the EV preparations were mixed with 250 μl PBS staining buffer containing 2.5 µg/ml DAPI (4’, 6-diamidino-2-phenylindole; Thermo Fisher Scientific, USA)) and kept at 4 °C for 30 min. The EVs were analyzed and sorted on the MoFlo Astrios cell sorter (Beckman Coulter) equipped with an EQ module specifically developed to detect nanoparticles and with 488 nm and 561 nm lasers at 200 mW. The calibration of the machine was carried out using FITC-labelled Megamix-Plus SSC beads from BioCytex (Fig. S1). The sheath-liquid 0.9% NaCl (Revol, France) was filtered through a 0.04 μm filter. The analysis was performed using the side-scattered (SSC) light parameter of laser 561, with threshold set to 0.012% in order to have maximum of 300 events per second. An M2 mask was added in front of the forward-scattered (FSC) light. All SSC and FSC parameters are viewed in logarithmic mode.

To count the EVs, we used Trucount™ Tubes (BD Biosciences, San Diego, CA), which contain a defined number of fluorescently labeled beads and have been specifically designed for reproducible counting of various biological nanoparticles, including EVs [54,55]. For quantification, we added the same volume (300 µl) of EV preparations into the tubes that contained the constant number of beads. The EV number was calculated using the following formula: EV_total_ = (EV count/bead count) × total number of beads in the Trucount™ Tube. In each case, the samples were passed through the flow cytometer’s detector until 2000 beads were recorded.

### Cell cycle synchronization

*S. islandicus* cells were synchronized as previously described for *S. acidocaldarius* [56,57]. For western blot, around 1×10^9^ cells were collected from the synchronized cell cultures. To isolate the EVs, 20 ml of cell culture at the indicated time points were collected and the EVs were isolated as described above.

### Cell cycle and cell size analysis by flow cytometry

The cell cycle of synchronized cells and the cell sizes of knockdown and over-expression strains were analyzed by flow cytometry. Around 0.6 × 10^8^ cells were collected for flow cytometry analysis. Briefly, cells at indicated time points were pelleted at 6,000 rpm for 5 min, resuspended in 300 µl of PBS, and then 700 µl of cool ethanol were added for at least 12 h to fix the cells. The fixed cells were then pelleted at 2,800 rpm for 20 min and washed with 1 ml of PBS. Finally, the cells were pelleted and resuspended in 80 µl of staining buffer containing 40 µg/ml propidium iodide (PI). After staining for 30 min, the DNA content or cell sizes were analyzed using the ImageStreamX MarkII Quantitative imaging analysis flow cytometry (Merck Millipore, Germany), which was calibrated with non-labeled beads with a diameter of 2 µm. The data from analysis of at least 100,000 cells was collected from each sample and analyzed with the IDEAS data analysis software.

### Live/Dead staining and fluorescence microscopy analysis

Live/Dead staining was carried out using the LIVE/DEAD *Bac*Light™ Bacterial Viability Kit (Invitrogen, US) [58,59] according to the supplier’s protocols. Specifically, around 0.6 × 10^8^ cells from the cell cultures at indicated time points were pelleted at 6,000 rpm for 5 min and resuspended in 50 µl of the M (mineral salts) solution. Then, the cells were mixed with 50 µl of the 2X stock solution of the LIVE/DEAD *Bac*Light staining regent mixture giving the final concentrations of SYTO 9 and propidium iodide of 6 µM and 30 µM, respectively. The samples were incubated at room temperature in the dark for 15 min, then the excess of dyes was removed by centrifugation at 6,000 for 5 min. The cells were resuspended in 80 µl of the M solution and 5 µl were used for fluorescence microscopy observation under the Leica TCS SP8 confocal microscope (Leica, Germany). The data was analyzed using the Leica Application Suite X imaging and analysis software (Leica, Germany).

For DAPI staining of the cells, around 0.6 × 10^8^ cells from the cell cultures were pelleted at 6,000 rpm for 5 min and resuspended in 80 µl of PBS staining buffer containing 9 µM (2.5 µg/ml) DAPI. For DAPI staining of the EVs, 50 µl of EV preparations were mixed with the 2× DAPI stock solution, giving a final concentration of DAPI of 9 µM (2.5 µg/ml). After staining for 30 min, 5 µl of DAPI-stained samples were used for fluorescence microscopy observation under the Leica TCS SP8 confocal microscope (Leica, Germany).

### Mass spectrometry and data analysis

The total protein content (i.e., membrane-associated and soluble proteins) of the Sis/pSeSD cells and highly purified (see above) EVs were analyzed by tandem liquid chromatography–tandem mass spectrometry (LC-MS/MS). The EVs and cells were snap-frozen in liquid nitrogen, lyophilized and re-suspended in 100 µl of lysis buffer including 8 M Guanidine HCl (GuHCl), 5 mM Tris(2-carboxyethyl)phosphine (TCEP) and 20 mM 2-chloro-acetamide (CAA). After kept at 95°C for 5 min, 900 µl of 50 mM Tris-HCl (pH 8.0) were added to the samples to dilute GuHCl to a concentration of under 1M. Then a mixture of 500 ng of LysC/Trypsin was added to the samples and kept at 37°C overnight for digestion of the proteins. The reaction was stopped by addition of 1% formic acid. Peptides were desalted using Sep-Pac C18 Cartridges (Waters, USA), following the manufacturer’s instructions. The purified peptides were concentrated to near dryness, re-suspended in 20 µl of 0.1% formic acid and analyzed by Nano LC-MS/MS at the Proteomics Platform of Institut Pasteur (Paris, France) using an EASY-nLC 1200 system (peptides were loaded and separated on a 30 cm long home-made C18 column; Thermo Fisher Scientific, Waltham, MA, USA) coupled to a Q Exactive Plus system (Thermo Fisher Scientific) tuned to the DDA mode. Peptide masses were searched against annotated *S. islandicus* REY15A proteins using Andromeda with the MaxQuant ver. 1.3.0.543 244 software, and additionally with the X!Tandem search engine. Identified proteins were functionally annotated against the archaeal clusters of orthologous groups (arCOG) database [60].

### DNA isolation from EVs and sequencing

For DNA extraction, EVs were collected during the exponential growth phase (24 h) and purified from 2 L of Sis/pSeSD culture, as described above. To remove the traces of extravesicular nucleic acids, EVs were incubated with DNase I (final concentration 15 U/ml) and RNase (final concentration of 100 μg/ml), in the presence of MgCl_2_ (final concentration of 10 mM), at 37°C for 30 min, followed by addition of EDTA (final concentration of 20 mM). EVs were disrupted by proteinase K (final concentration of 100 μg/ml) and SDS (final concentration of 0.5%) treatment at 55°C for 30 min. The DNA was extracted by standard phenol/chloroform procedure, precipitated with 0.3 M sodium acetate (pH 5.3) and isopropanol. The resultant pellet was resuspended in DNase/RNase-free water and used for sequencing. Sequencing libraries were prepared from 100 ng of DNA with the TruSeq DNA PCR-Free library Prep Kit from Illumina and sequenced on Illumina MiSeq platform with 150-bp paired-end read lengths (Institut Pasteur, France). Raw sequence reads were processed with Trimmomatic v.0.3.6 and mapped to the reference genomes of REY15A and pSeSD plasmid using Bowtie2 [61] with default parameters and analyzed with Sequana [62].

### Heterotrophic growth assay

EVs were isolated from 20 L of Sis/pSeSD cell culture and purified by ultracentrifugation in sucrose gradient as described above. The EVs were resuspended in 18 ml of the M (mineral salts) solution. The final concentration of the EV preparation was 49.6 μg/ml, based on the total protein amount. *S. islandicus* REY15A cells were cultured in MTSV medium and collected when they reached the early logarithmic phase (OD_600_=0.2). The cells were washed 6 times with the M solution by centrifugation at 7,000 rpm for 10 min to remove the traces of sucrose (S) and tryptone (T). The washed cells were inoculated into 10 ml of MV, MSV and MTSV medium as the control groups, to the initial OD_600_ of 0.05. Experimental groups were each supplemented with 1.5 ml of the EV preparations containing different concentration of EVs (24.8 μg for group I, 49.6 μg for group II and 74.4 μg for group III). The experiment was repeated three times.

### Construction of the CRISPR type III-B-based RNA interference plasmids and RNA interference

The CRISPR type III-B-based RNA interference plasmids were constructed according to the methods described previously [63,64]. For RNA interference, 40-nt protospacers matching the genes of interest were selected from the anti-sense strand of the corresponding genes downstream of the GAAAG, CAGAG or AAAG (5’-3’) sequences and cloned into the genome-editing plasmid pGE [63]. The spacers selected and used in this study are listed in Table S4. Spacer fragments were generated by annealing the corresponding complementary oligonucleotides and inserted into pGE at the BspMI restriction site. Plasmid pGE was introduced into E233S cells by electroporation and transformants were selected on MSCV plates without uracil. RNA interference was induced by arabinose (0.2% wt/vol final concentration). For enumeration with flow cytometry, the EVs were collected from 50 ml of cell cultures during the exponential growth phase of different knockdown strains (24 h), as described above. The flow cytometry measurements were done in triplicate (Fig. S16).

### RNA preparation and quantitative reverse-transcription PCR (RT-qPCR)

Cells from RNA interference (knockdown) strains were collected for RNA extraction after 24 h post induction. Total RNAs were extracted using TRI Regent® (SIGMA-Aldrich, USA). The concentrations of the total RNAs were estimated using the Eppendorf BioSpectrometer® basic (Eppendorf AG, Germany). The quality of the RNA preparations was further checked by agarose gel electrophoresis.

First-strand cDNAs were synthesized from the total RNAs according to the protocol of Maxima First Strand cDNA Synthesis Kit for RT-qPCR with dsDNase (Thermo Scientific, USA). Shortly, 2 μg of RNA were treated with the dsDNAase at 37°C and then the reverse transcription reaction was carried out by incubation for 10 min at 25°C followed by 30 min at 50°C, and finally terminated by heating at 85°C for 5 min. The resulting cDNA preparations were used to evaluate the mRNA levels of the targeted genes by qPCR (2ng of cDNA were used as the template), using Luna® Universal qPCR Master Mix (New England Biolabs, USA) and gene-specific primers (Table S5). qPCR was performed in an Eppendorf MasterCycler RealPlex^4^ (Eppendorf AG, Germany) with the following steps: denaturing at 95°C for 2 min, 40 cycles of 95°C 15 s, 55°C 15 s and 68°C 20 s. Relative amounts of mRNAs were calculated using the comparative Ct method with 16S rRNA as the reference [65]. Three independent biological experiments and three technical replicates were carried out for RT-qPCR.

### Overexpression of ESCRT proteins

Plasmids expressing different ESCRT machinery components and their mutants were described previously [38]. Briefly, cells harboring the plasmids were first inoculated into 30 ml of the MTSV medium and when the OD_600_ reached ∼0.6-0.8, they were transferred into the ATV medium containing 0.2% (wt/vol) arabinose with an initial OD_600_ of 0.05 to induce protein expression. All plasmids are listed in Table S3.

### EV-mediated gene transfer

EVs isolated and purified from 6 L of exponentially growing Sis/pSeSD culture (24 h) were used for gene transfer experiments as described previously [66], with some modification. Briefly, E233S cells were grown in 30 ml of MTSVU medium until optical density reached 0.2 and then harvested by centrifugation (7,000 rpm for 10 min at room temperature). The cell pellet was washed 6 times with 30 ml of 0.7% (wt/vol) sucrose solution to remove the uracil and then resuspended in 30 ml of MCSV. One ml of EV preparation (76 μg/ml based on the total protein amount) or PBS (control) were added to 5 ml of E233S cell culture. The cells were incubated at 75°C with shaking (145 rpm). After 3 h, 5 h and 7 h of incubation 100 μl of each sample were collected and serial dilutions were spread on the pre-warmed MCSV plates and incubated at 75°C. After 10 days of incubation, single colonies were picked, inoculated into nuclease free water and 2 μl were used as template for PCR with plasmid-specific primers pSeSD-F and pSeSD-R (Table S5) to check for the presence of pSeSD plasmid.

### Spot test

Colonies formed on the MSCV plates were picked with a toothpick and resuspended in 100 μl of the M solution. Then, 10 μl of the cell suspension were spotted on the pre-warmed MSCV plates and incubated at 75°C for 3-5 days.

### Western blot

To verify the presence of ESCRT proteins in the EVs, the sucrose-gradient purified EV samples were run in 12% polyacrylamide gel using tris-glycine running buffer, then transferred onto PVDF membrane. ESCRT proteins were detected using antibodies against ESCRT-III, ESCRT-III-1 and ESCRT-III-2 (HuaAn Biotechnology Co., Hangzhou, Zhejiang, China), as described previously [38]. The goat anti-rabbit (Thermo Fisher Scientific, USA) secondary antibodies coupled with peroxidase were used as secondary antibodies. The specific bands were detected by chemoluminescence using ECL prime western blotting detection reagents (Amersham) according to the manufacturer’s instructions. Proteins purified from *E. coli* BL21-CodonPlus(DE3)-RIL (Agilent Technologies) were used as the positive controls.Materials and Methods are available in the Supplementary Information.

## RESULTS AND DISCUSSION

### EV production and purification

To study the composition and function of archaeal EVs, and to investigate the role of ESCRT in their biogenesis, we established a procedure for purification and quantification of EVs from *Sulfolobus islandicus* REY15A and *Saccharolobus solfataricus* PH1 cells (Sis-EVs and Sso-EVs, respectively), and compared the EV production at different growth stages in both strains. Consistent with the observations made for EVs isolated from other *Sulfolobus* species [49], Sis-EVs and Sso-EVs were visibly coated with the proteinaceous surface (S-)layer (Fig. 1a) typical of *Sulfolobus* cells [67,68] and displayed considerable variation in shape and diameter. The median diameters of Sis-EVs and Sso-EVs were 176.54 nm and 185.85 nm, respectively, with the Sso-EVs being slightly more variable in size (Fig. 1b). The EVs were collected at different stages of cell growth (Fig. 1c) and could be reproducibly quantified by flow cytometry using tubes containing a calibrated number of fluorescent beads (Fig. 1d and Fig. S1). With a notable exception of the 12-hour time point, EV production by *S. islandicus* and *S. solfataricus* followed a similar pattern: EV titer increased throughout the growth of the cells (Fig. 1c). Given the similarities in EV production patterns in *S. islandicus* and *S. solfataricus*, for all subsequent experiments, we focused on EVs from *S. islandicus*, for which more advanced genetic tools are available [69]. We next tested whether there is a link between Sis-EV production and increase in the fraction of the dead cells in the growing *S. islandicus* population by performing the live/dead staining at different time points (see Materials and Methods). From 12 to 60 h (early exponential to stationary phase) the ratio of dead cells remained at around 1% and only when the cells progressed into the ‘death’ phase, the ratio of dead cells increased sharply, with around 30% of dead cells at 72 h and more than 90% of dead cells at 84 h (Fig. S1). These results suggest that Sis-EVs production is not a consequence of cell death. To verify that the EV preparations were devoid of cellular contaminants, we performed the following procedures: (i) flow cytometry profiles of the EV samples were compared with those containing *Sulfolobus* cells (Fig. S2); (ii) purified EV preparations were subjected to semi-quantitative transmission electron microscopy analysis (Fig. S3a); (iii) EV preparations were also analyzed by fluorescence microscopy (Fig. S3b); and (iv) EV preparations were plated on solid medium supporting the growth of *Sulfolobus* cells (Fig. S4). None of these procedures revealed the presence of *Sulfolobus* cells, viable or otherwise, in the EV preparations.

**Fig. 1.**
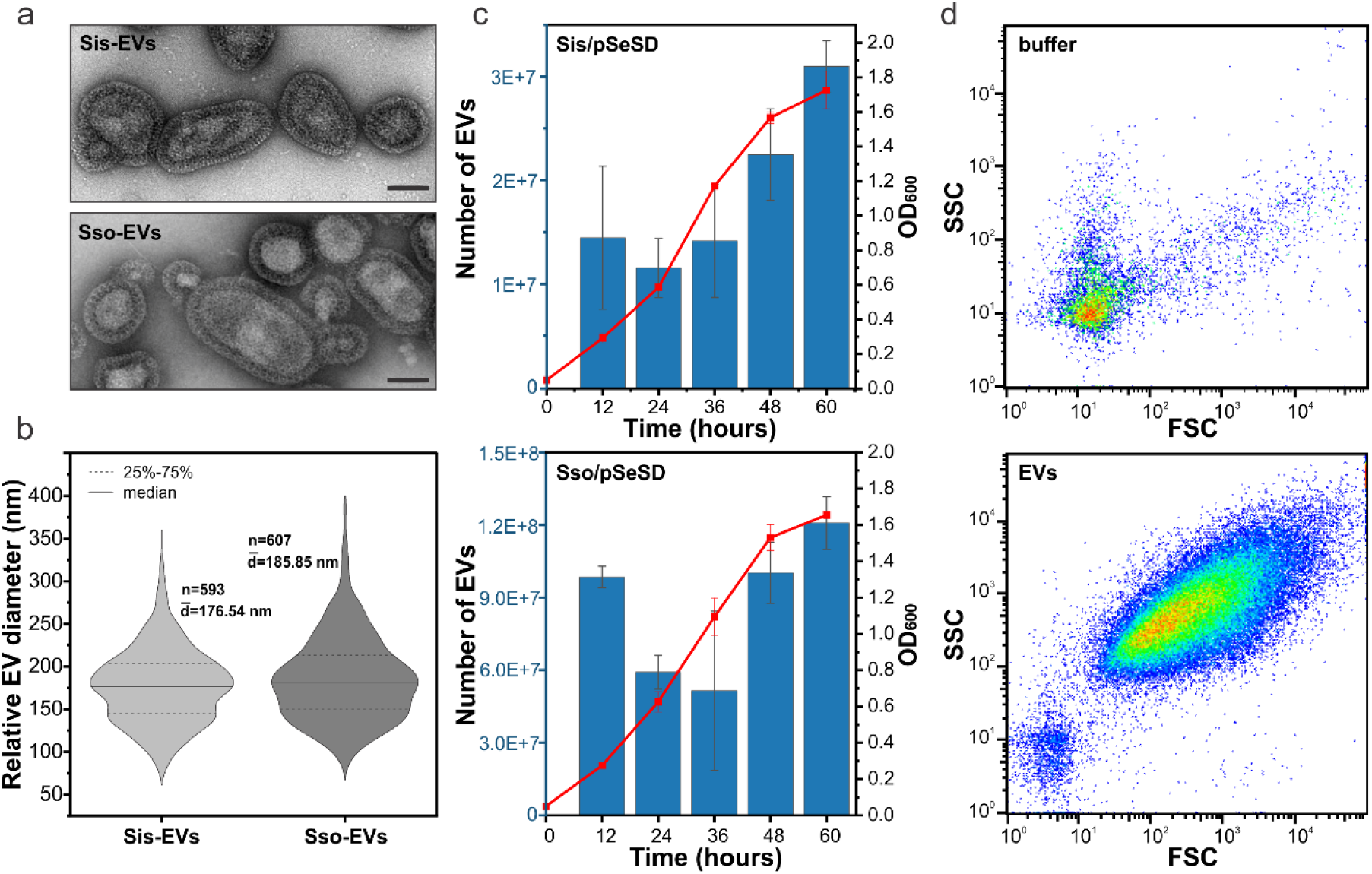
Characterization of the *Sulfolobus islandicus* EVs. **a** Transmission electron micrographs of negatively stained Sis-EVs (top) and Sso-EVs (bottom). Scale bars, 100 nm. **b** Violin plots showing the size distributions of Sis-EVs (n=593) and Sso-EVs (n=607). The width of the distribution corresponds to the frequency of occurrence. **c** Growth curves of *S. islandicus* E233S and *S. solfataricus* PH1-16 harboring the vector pSeSD, and quantification of EVs released at indicated time points. Error bars represent standard deviation from three independent experiments. **d** Quantification of Sis-EVs by flow cytometry. Top panel shows the control with buffer only. SSC, side scattering; FSC, forward scattering.

### Protein content of Sis-EVs

EVs are known to carry diverse cargo, including proteins [2-5]. Proteomics analysis of Sis-EVs and *S. islandicus* cells led to the identification of 413 and 1035 proteins, respectively (Supplementary Data 1). The number of proteins detected in Sis-EVs is considerably higher than that reported previously for EVs from *S. acidocaldarius, S. solfataricus* and *S. tokodaii* [49], possibly due to improved sensitivity of mass-spectrometry over the past decade. Notably, recent studies on the proteomics of EVs produced by diverse bacteria [70-73] as well as halophilic archaea [18] report the presence of hundreds of proteins in each type of EVs, consistent with our results. For instance, it has been shown that EVs produced by halophilic archaeon *Halorubrum lacusprofundi* contain 447 different proteins [18].

All but one functional protein categories found in *S. islandicus* proteome, as defined using the archaeal clusters of orthologous groups (arCOG; Table S1) [60], were represented in the Sis-EVs (Fig. 2a). Proteins of the arCOG category X (Mobilome: prophages, transposons) were not found in the Sis-EVs, likely due to the fact that only few proteins of this functional category are expressed in *S. islandicus* under normal growth conditions [74,75]. The fractions of proteins of the categories J (Translation, ribosomal structure and biogenesis), K (Transcription), V (Defense mechanisms), H (Coenzyme transport and metabolism) and I (Lipid transport and metabolism) were more than twice smaller compared to their corresponding fractions in the total cellular proteome. By contrast, arCOG categories D (Cell cycle control, cell division, chromosome partitioning), N (Cell motility), O (Posttranslational modification, protein turnover, chaperones), U (Intracellular trafficking, secretion, and vesicular transport), C (Energy production and conversion), P (Inorganic ion transport and metabolism) and S (Function unknown) were enriched in Sis-EVs compared to the total cellular proteome (Fig. 2a). For instance, the D category proteins constitute only 0.6% of the total *S. islandicus* proteome, whereas in Sis-EVs, these proteins correspond to 1.7% of proteins (2.9-fold increase). There is also notable enrichment in Sis-EVs of proteins with predicted transmembrane domains compared to the cellular proteome (20% vs 4%; Fig. 2b). Of the top 100 most abundant proteins in the EVs, 65 have predicted transmembrane domains, whereas there are no such proteins among the top 100 most abundant cellular proteins. Thus, although Sis-EVs incorporate a considerable fraction of the total

**Fig. 2.**
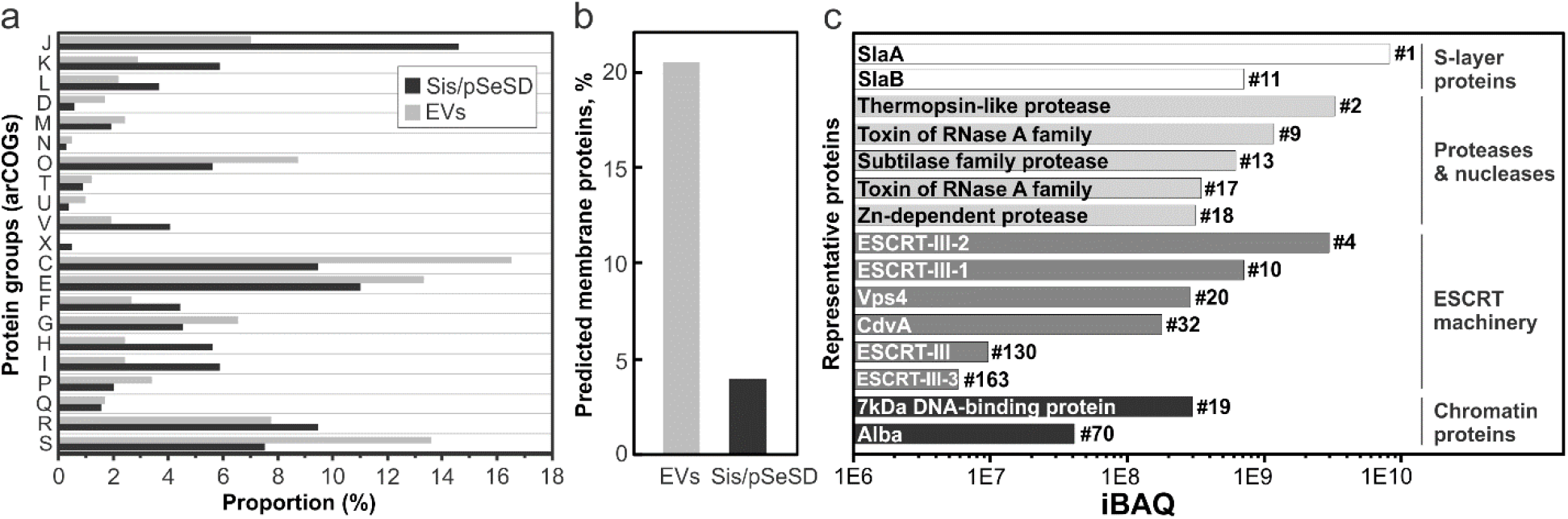
Analysis of the Sis-EV protein content. The EVs were collected during the exponential growth phase (24 h) of Sis/pSeSD, purified on sucrose gradient, treated with DNase I and subjected to mass spectrometry analysis. **a** Functional classification of proteins identified in highly purified Sis-EVs using archaeal clusters of orthologous groups (arCOGs). arCOG categories are indicated with capital Roman letters, with the annotation provided in Table S1. **b** Fraction of proteins with predicted transmembrane domains in Sis-EV and cellular (Sis/pSeSD) proteomes. **c** Label-free intensity-based absolute quantification (iBAQ) of selected Sis-EV proteins (the corresponding functional categories are indicated on the right). Numbers next to each bar indicate the abundance rank.

*S. islandicus* proteome, there is a strong enrichment for certain functional categories and, in particular, for membrane proteins. Presence in the Sis-EVs of proteins from nearly all functional categories suggests that many of these proteins are incorporated non-selectively, by entrapment of the cytosolic and membrane contents. It is possible that not all of the proteins are present within each Sis-EV, but are rather distributed across the EV population.

*Sulfolobus* EVs have been previously shown to carry toxins, dubbed sulfolobicins, active against closely related *Sulfolobus* species [76,77]. However, homologs of these particular toxins are not encoded in the *S. islandicus* REY15A genome. Nevertheless, nearly one third of proteins in the O category in Sis-EVs corresponded to diverse proteases and nucleases. Notably, we also detected two putative toxins (WP_014512538 and WP_014512541) of the RNase A family and several hydrolases of diverse specificities (Fig. 2c; Supplementary Data 1). This finding suggested that deployment of the Sis-EV payload could be toxic to recipient cells lacking necessary immunity. Incubation of Sis-EVs with *Sulfolobus* cells for 3 h indeed resulted in modest, albeit significant, decrease in colony forming units for *S. solfataricus* cells, but not for the more distantly related *S. acidocaldarius* or *S. shibatae* (Fig. S5a). However, the inhibitory effect of Sis-EVs on *S. solfataricus* was temporary and was lifted when the incubation was prolonged to 5 h (Fig. S5b). Thus, Sis-EVs do not appear to participate in intermicrobial conflicts, at least, not among the tested *Sulfolobus* species.

Sis-EVs contained all six components of the *Sulfolobus* ESCRT machinery (arCOG category D; Fig. 2c), consistent with the possibility that ESCRT machinery is involved in EV biogenesis [49]. Label-free intensity-based absolute quantification (iBAQ) [78] of protein abundances showed that two of the ESCRT components, ESCRT-III-2 and ESCRT-III-1, were among the top-10 most abundant proteins in Sis-EVs (Fig. 2c). Western blot analysis has confirmed that both proteins were present and strongly enriched in Sis-EVs (Fig. S6). As expected, both S-layer proteins were found in the EVs, with SlaA being the most abundant protein in Sis-EVs (Fig. 2c).

### Sis-EVs carry plasmid and genomic DNA

Sis-EVs carried the chromatin proteins Sac7d/Sso7d and Alba (Fig. 2b), responsible for compaction of the *Sulfolobus* chromosome [79,80], suggesting that Sis-EVs contain DNA. Indeed, DNA has been previously observed in EVs from halophilic archaea and *Thermococcus* spp. (both in the phylum Euryarchaeota) [18,19,21,81] but never reported in *Sulfolobus* EVs. To test if Sis-EVs carry DNA, purified EVs were treated with DNase I and stained with 4′,6-diamidino-2-phenylindole (DAPI). The DAPI-stained Sis-EVs could be detected by both flow cytometry (Fig. 3a) and fluorescence microscopy (Fig. 3b), consistent with the presence of DNA. Notably, only 13.3% of EVs detected by the flow cytometry were DAPI-positive, whereas the majority of EVs were DAPI-negative, indicating a heterogeneity of the EV content.

**Fig. 3.**
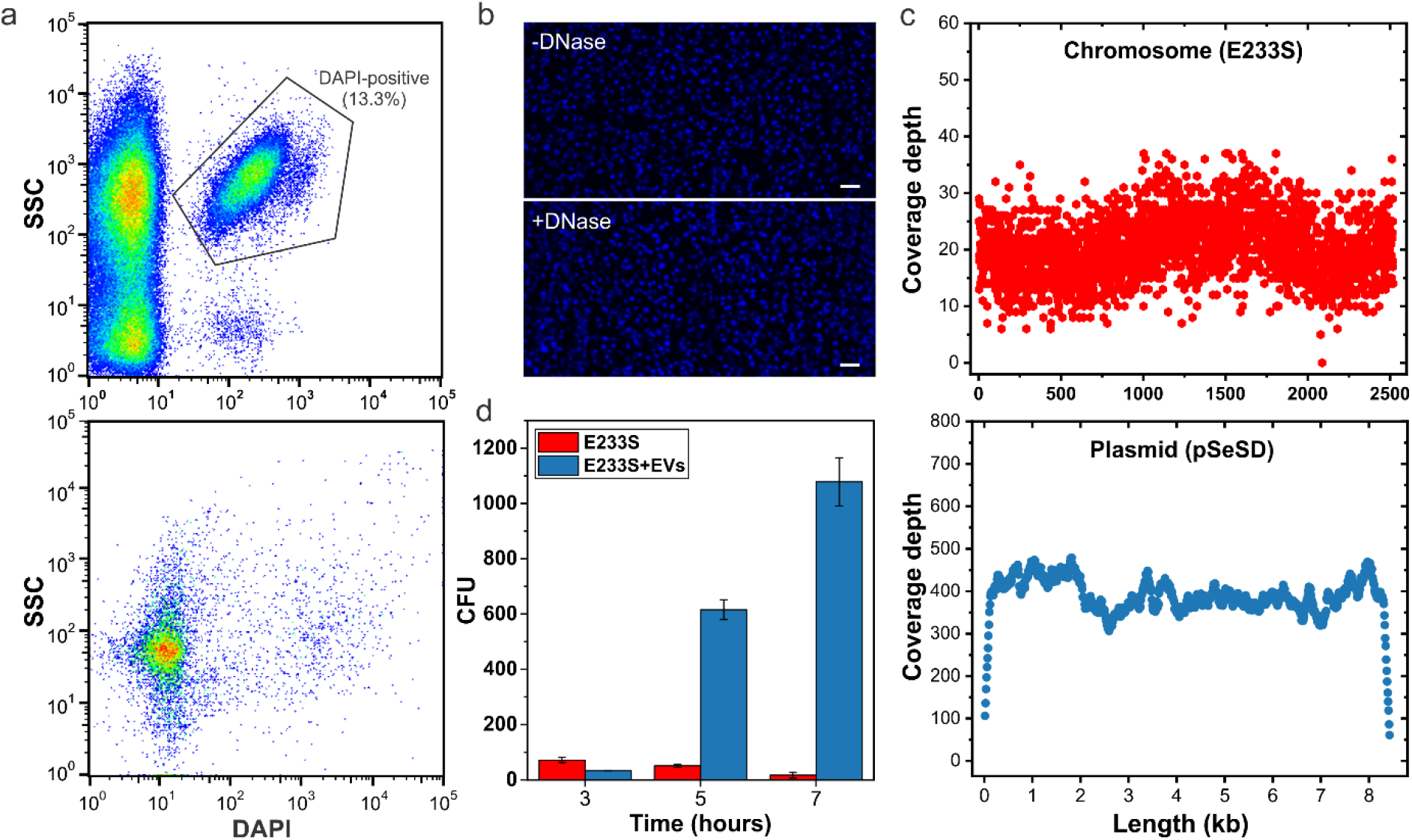
Sis-EVs promote gene transfer. **a** Analysis of DAPI-stained Sis-EVs isolated from Sis/pSeSD (upper panel) by flow cytometry. All the events are shown, with the selected region indicating the DAPI-positive EVs harboring DNA. Note that most EVs are DAPI-negative. Bottom panel shows the control with the DAPI-containing buffer only. SSC, side scattering. **b** Fluorescence micrographs of DAPI-stained Sis-EVs prior (top) and after (bottom) DNase I treatment. Bar, 2 μm. **c** Sequencing depth across the chromosomal DNA (E233S) and plasmid (pSeSD). Each dot represents sequencing depth at the indicated position of the corresponding replicon. **d** Gene/plasmid transfer by Sis-EVs. Sis-EVs were treated with DNase I and then mixed with E233S cells, incubated for 3, 5 and 7 h and plated on selective plates. In the control experiment, E233S cells were mixed with the equal volume of PBS buffer. The number of obtained colony forming units (CFU) is plotted on the y-axis. Error bars represent standard deviation from three independent experiments.

High-throughput sequencing of the DNA extracted from Sis-EVs yielded reads aligning to both the *S. islandicus* chromosome and the resident extrachromosomal plasmid pSeSD. Both replicons were covered throughout their respective lengths (Fig. 3c), but the median sequencing depth of the plasmid was 19 times higher than that of the chromosome (386× versus 20× coverage, respectively). This difference cannot be explained by the higher copy number of the plasmid (3-5 copies per 1 chromosome copy) [51]. It is most probable that, as in the case of the *Thermococcus* [82] and bacterial [8] EVs, overlapping genomic fragments of variable sizes, rather than complete chromosome, are packed into the Sis-EVs.

To test the biological relevance of DNA incorporation into Sis-EVs, we investigated the ability of Sis-EVs to transfer the plasmid-borne *pyrEF* locus into a plasmid-free auxotrophic strain E233S of *S. islandicus* carrying a chromosomal deletion in the *pyrEF* operon responsible for uracil synthesis (Fig. S7) [51]. To this end, Sis-EVs produced by the pSeSD-carrying strain were purified, treated with DNase I, incubated with recipient E233S cells in liquid culture and plated on solid medium devoid of uracil. In the presence of EVs, strain E233S formed over 1000-fold more colonies than the control E233S cells (Fig. 3d), whereas plating of EVs alone did not yield colonies on either rich or uracil-deficient medium (Fig. S8a), further confirming that there were no cells contaminating EV preparation. Approximately half of the colonies obtained after incubation with Sis-EVs carried the pSeSD plasmid (Fig. S8b). To test if the plasmid-devoid strains carry the *pyrEF* cassette elsewhere on the chromosome, we performed PCR with the *pyrF*-specific primers (Fig. S7). However, *pyrF* gene was present exclusively in the pSeSD-carrying strains (Fig. S8c), indicating that there was no ectopic *pyrF* integration. Next, we verified if the ability of the plasmid-lacking strains to grow in the absence of uracil was inheritable. To this end, the plasmid-containing and plasmid-deficient cells from initial colonies were resuspended in the selective medium and spotted on the solid medium lacking uracil (Fig. S8d). Only plasmid-containing strains could grow, suggesting that initial growth of the colonies in the absence of uracil was supported by the nutrients provided by the EVs, whereas transfer of such colonies into fresh medium arrested the cell growth, unless the cells contained pSeSD. Collectively, these results demonstrate that Sis-EVs carry DNA and act as vehicles for gene transfer. The exact mechanism of EV-mediated gene transfer into recipient cells remains unclear but, presumably, it involves fusion between the EV and cell membranes.

### Sis-EVs support heterotrophic growth of *Sulfolobus* cells

To further investigate if EVs can provide nutrients (other than uracil) to support heterotrophic cell growth, *Sulfolobus* cells were inoculated in media lacking nitrogen and/or carbon source. As expected, cells could not grow in the medium containing only mineral salts and a mix of vitamins (Fig. 4a), nor could they grow when only carbon source (sucrose) was added to this solution (Fig. 4b). Instead, slight decrease in optical density of the culture was observed, suggesting partial lysis. However, when either medium was supplemented with purified Sis-EVs, there was significant (two paired T-test, p <0.05), Sis-EV concentration-dependent increase in the optical density of *S. islandicus* culture, indicative of cell growth (Fig. 4). The same result was obtained with EVs isolated from Sis/pSeSD-CdvA (see below). These results strongly suggest that EVs can serve as a source of both carbon and nitrogen, and hence play an important role in nutrient cycling in extreme environments. Similarly, it has been shown that EVs produced by cyanobacteria can support the growth of heterotrophic bacteria by serving as a carbon source [7].

**Fig. 4.**
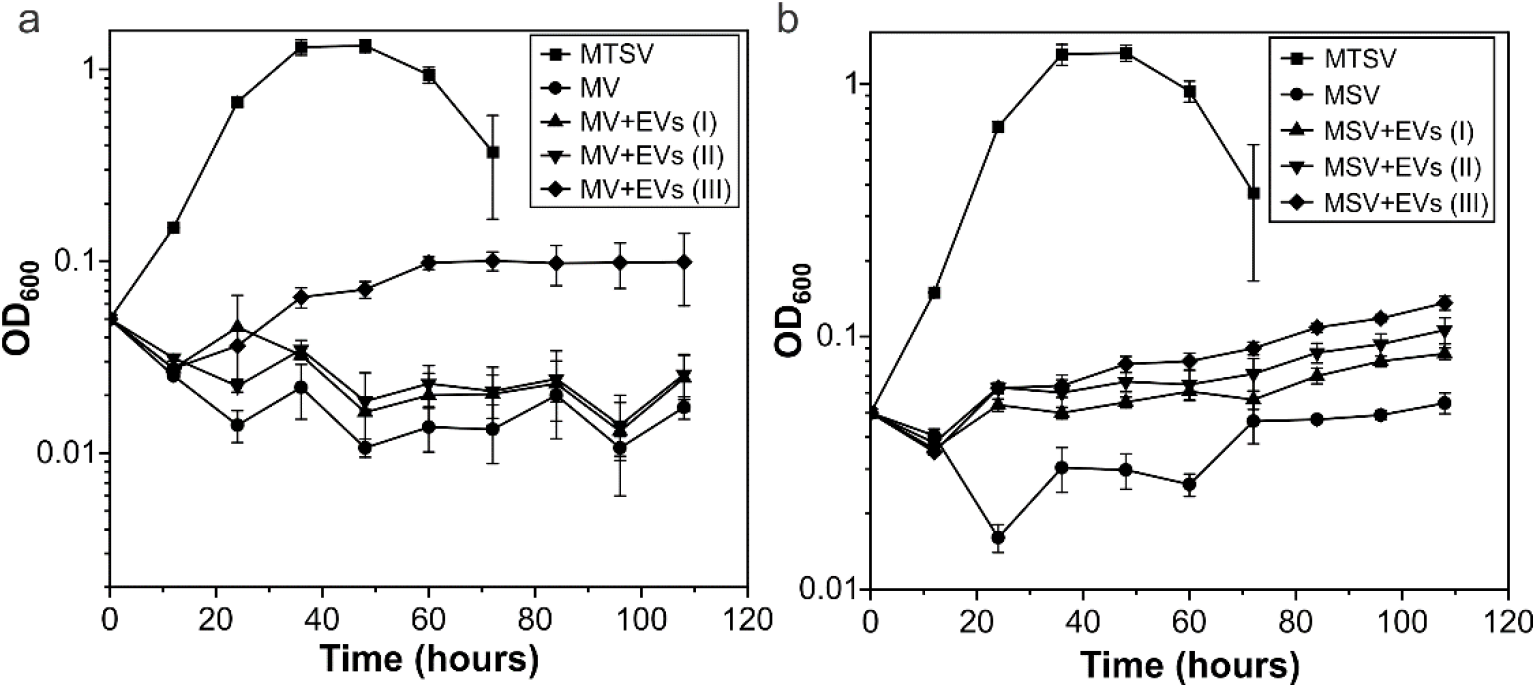
Sis-EVs support heterotrophic growth of *Sulfolobus* cells. **a** Growth curves of *S. islandicus* REY15A in MV (M: mineral salts, V: vitamin mix) solution lacking organic carbon and nitrogen sources; MV solution supplemented with 1.5 ml preparations containing different amounts of EVs (MV+EVs): 24.8 μg for I, 49.6 μg for II, and 74.4 μg for III; and rich medium (MTSV), which in addition to MV solution contains 0.2% (wt/vol) sucrose (S) and 0.2% (wt/vol) tryptone (T). There was ssignificant increase (two paired t-test, p <0.05) in optical density (OD_600_) of REY15A cells in the MV solution supplemented with EVs (III), as compared to the control culture lacking EVs. **b** Growth curves of *S. islandicus* REY15A in MSV (M: mineral salts, S: sucrose, V: vitamin mix) medium lacking organic nitrogen source; MSV medium supplemented with different concentrations of EVs (MSV+EVs), and rich MTSV medium. There was significant increase (two paired t-test, p <0.05) in optical density (OD_600_) of REY15A cells in the MSV medium supplemented with EVs, as compared to the control culture lacking EVs. Error bars represent standard deviations from three independent experiments.

To the best of our knowledge, production of EVs by different Sulfolobales strains or by any other archaeal strain has been reported only under laboratory cultivation conditions. To verify whether EVs are also produced in the environment, we analyzed a previously collected archaea-dominated hot spring sample [53] for the presence of EVs. The contents of the sample were directly concentrated by ultracentrifugation without prior cultivation in the laboratory and visualized by TEM. We observed multiple S-layer-coated EVs closely resembling those produced by Sulfolobales species (Fig. S9). The diameter of the observed EVs varied between 77 and 182 nm, which is considerably smaller than the size of the smallest known archaea, i.e., *Nanoarchaea* spp. with the diameter of ∼400 nm [83,84], confirming that these are subcellular structures. These results strongly suggest that Sis-EVs are not laboratory artifacts and are environmentally and biologically relevant.

### Sis-EVs biogenesis is ESCRT-dependent

Sis-EV biogenesis occurs through budding from the cytoplasmic membrane (Fig. 5a) and ESCRT system is a prime suspect implicated in membrane constriction and scission. To investigate the involvement of ESCRT machinery in EV biogenesis, we constructed a collection of knockdown strains in which expression of each of the six ESCRT machinery components was depleted by the endogenous type III-B CRISPR-Cas system of *S. islandicus*. The utility of this strategy for gene knockdown has been recently demonstrated in *Sulfolobus* [63,68,85]. Quantitative reverse transcription PCR (RT-qPCR) analysis has shown that whereas expression of *escrt-III* was decreased by ∼30%, expression of all other genes was down by 60-70% (Fig. 5b). Western blot analysis of *escrt-III-1* and *escrt-III-2* knockdown strains has shown that the levels of the corresponding proteins have been decreased by 99% and 60%, respectively (Fig. S6b). It has been previously shown that *escrt-III* and *vps4* are expressed from a bicistronic operon [44,86] (Fig. S10a). Thus, we verified whether CRISPR targeting of the *escrt-III* gene has a polar effect on the expression of the *vps4*. There was no significant difference in the *vps4* transcript levels between the *escrt-III* knockdown and control cells (Fig. S10b). This is consistent with the previous results showing that cleavage of a transcript by type III-B CRISPR system in *S. islandicus* REY15A occurs within 20 bp of the CRISPR spacer targeting [64]; that is, the fragment of the transcript encoding Vps4 is unaffected by the cleavage within the ESCRT-III-encoding region (Fig. S10a).

**Fig. 5.**
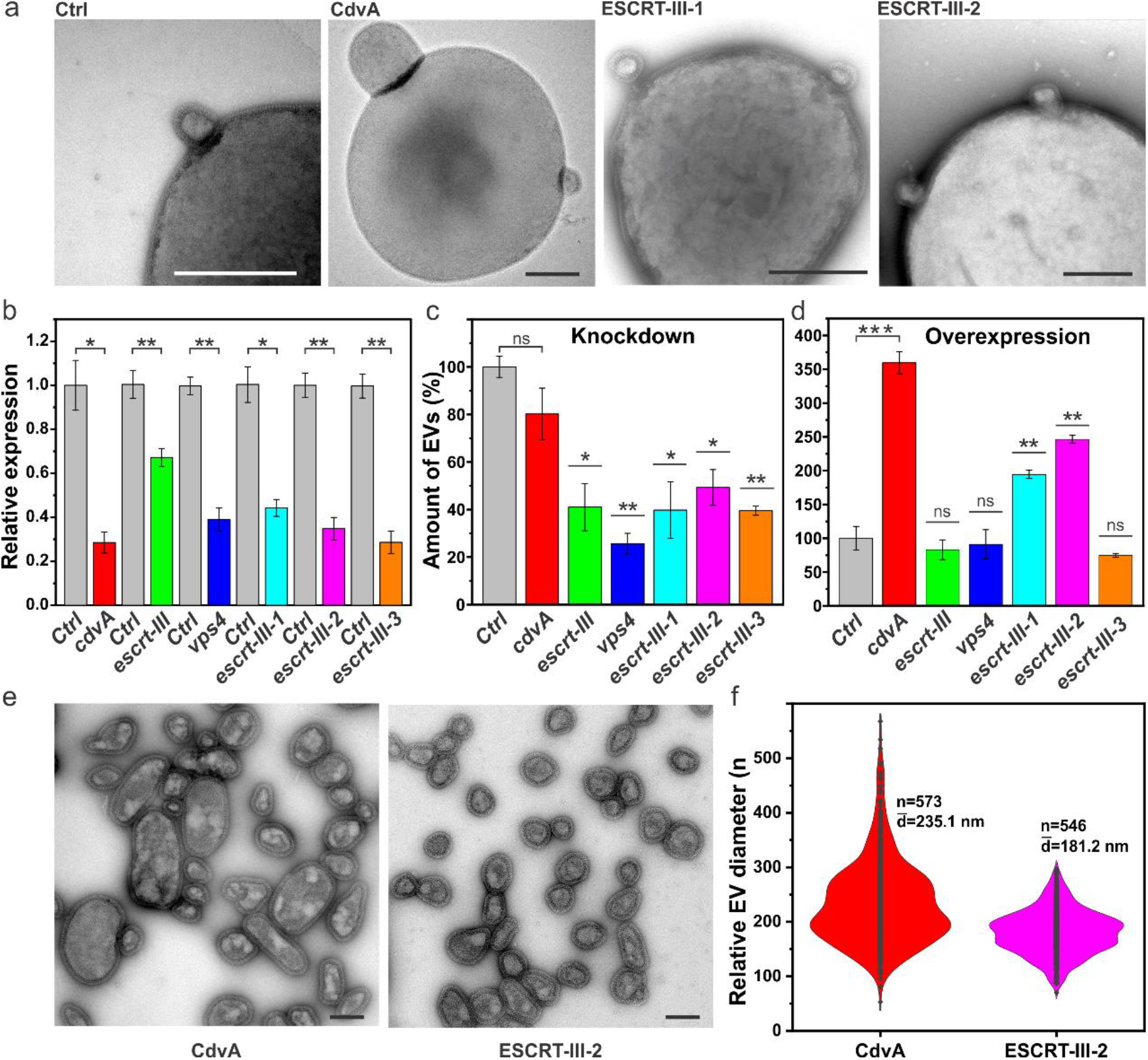
ESCRT-dependent biogenesis of Sis-EVs. **a** Representative transmission electron micrographs showing EV budding from *S. islandicus* strains overexpressing indicated proteins. Ctrl: control E233S cells carrying empty vector pSeSD. Bars, 400 nm. **b** RT-qPCR analysis of the RNA interference efficiency. Stars indicate the significance levels based on the paired two-tailed t-test. The p-values are 0.01512, 0.00514, 0.00737, 0.0146, 0.00883, 0.00733, respectively. Error bars represent standard deviation from three independent experiments. **c** Quantification of Sis-EVs released from strains in which different ESCRT machinery components were depleted by CRISPR targeting. Stars indicate the significance levels based on the paired two-tailed t-test. The p-values are 0.01047, 0.00316, 0.02337, 0.01763 and 0.00177. ns, non-significant. Error bars represent standard deviation from three independent experiments. **d** Quantification of Sis-EVs released from strains overexpressing indicated ESCRT machinery components. Stars indicated the significance levels based on the paired two-tailed t-test. The p-values are 0.001, 0.0094 and 0.00435. ns, non-significant. Error bars represent standard deviation from three independent experiments. **e** Representative transmission electron micrographs of negatively stained Sis-EVs isolated from cells overexpressing CdvA and ESCRT-III-2. Bars, 200 nm. **f** Violin plots showing the size distributions of Sis-EVs isolated from cells overexpressing CdvA (n=573) and ESCRT-III-2 (n=546). The width of the distribution corresponds to the frequency of occurrence.

Knockdown strains of *cdvA, vps4, escrt-III* and *escrt-III-1* displayed considerable cell growth defects, whereas those of *escrt-III-2* and *escrt-III-3* showed nearly normal cell growth (Fig. S11a). The depletion of *cdvA, vps4, escrt-III* and *escrt-III-1* transcripts by CRISPR targeting resulted in obvious cell division defects (Fig. S11b, c), yielding cells 2-3 times larger than the control cells, and in slight increase (<7%) in the fraction of dead cells in the corresponding populations (Fig. S11d). Consistent with the growth dynamics (Fig. S11a), cell size of the *escrt-III-2* and *escrt-III-3* knockdown strains was similar to that of the control cells (Fig. S11b, c). The lack of growth retardation for the *escrt-III-2* knockdown strain is somewhat unexpected, given that all previous attempts to delete this gene in *S. islandicus* were unsuccessful, whereas *escrt-III-3* is known to be non-essential for normal growth [38,87]. Notably, under the growth conditions used in this study, the expression of *escrt-III-3* was much lower than that of all other cell division genes (Fig. S12). By contrast, the transcription level of *escrt-III-2* was six and three times higher than those of *escrt-III* and *escrt-III-1*, respectively (Fig. S12). Thus, even with 60-70% decrease in transcript levels due to CRISPR targeting, the total level of *escrt-III-2* transcripts would be comparable to that of other ESCRT-III homologs. Presumably, these levels are sufficient for normal growth of *S. islandicus*. Notably, *escrt-III-2* is not essential in *S. acidocaldarius* [40]. Thus, we do not exclude the possibility that *escrt-III-2* is also not strictly required for the growth of *S. islandicus* and that previous attempts to delete this gene were hindered by other factors.

Quantification of Sis-EVs produced by the knockdown strains has shown that whereas depletion of *cdvA* had no significant effect on Sis-EV titer, all other knockdown strains, including the non-essential *escrt-III-3*, produced significantly less EVs compared to the control strain (Fig. 5c). The *vps4* knockdown strain displayed the strongest effect, with EV production being decreased by over 70%. Notably, it is possible that different ESCRT-III homologs can partially complement each other during EV biogenesis. Interestingly, overexpression of the ESCRT-III-1 and ESCRT-III-2 from a plasmid resulted in 200-250% increase in vesiculation (Fig. 5d) consistent with their role in Sis-EV budding. Unexpectedly, overexpression of CdvA resulted in hypervesiculation phenotype (Fig. 5d). However, the same effect was also observed when CdvA lacking the C-terminal domain (CdvAΔC) responsible for interaction with ESCRT-III [43,44] was overexpressed (Fig. S13a), suggesting that excessive binding of CdvA to the membrane [44] precipitates the observed phenomenon. Overexpression of *cdvA* and *cdvAΔC* yielded cells with up to 2-5 fold larger diameters (Fig. S13b). By contrast, overexpression of ESCRT-III-1 and ESCRT-III-2 had no effect on cell size or cell viability (Fig. S13c). Taken together, the overexpression and knockdown results show that there is no apparent link between the cell size and EV biogenesis.

Budding of EVs from the control and overexpression strains was observed directly by TEM (Fig. 5a). Notably, EVs produced by the CdvA overexpression strain were considerably larger than those from the control (Fig. 1d) and ESCRT-III-2 overexpression (Fig. 5e) strains, with an average diameter of 235 nm versus 177 and 181 nm, respectively (Fig. 5f). To exclude the possibility that the large EVs produced by the CdvA overexpression strain represent small cells, the EV-containing supernatant was filtered through 0.45 μm filter and plated on the solid medium. No colonies were formed (Fig. S4b). Our results strongly suggest that EV budding in *Sulfolobus* is dependent on the ESCRT machinery, including Vps4 ATPase and the ESCRT-III ensemble, whereas CdvA appears to be dispensable for this process.

### Sis-EVs biogenesis is linked to cell division

The expression of ESCRT-III homologs in *S. acidocaldarius* is linked to the cell cycle [37,56]. To verify whether the same is true for *S. islandicus* and if EV biogenesis is linked to the cell cycle, we synchronized the *S. islandicus* culture by adapting a protocol previously used for *S. acidcaldarius* [56,57]. The cells were arrested at the G2 phase using acetic acid and could progress into cell division phase following the removal of the acid (see Materials and Methods for details). Analysis of the DNA content by flow cytometry has shown that the cells started transitioning from G2 into the cell division phase at 90 min following the removal of the acetic acid (Fig. 6a). Western blot analysis of the synchronized cells has shown that ESCRT-III, ESCRT-III-1 and ESCRT-III-2 proteins were undetectable during the G2 phase and became detectable at 90 min after the removal of acetic acid (Fig. 6b). Notably, however, whereas ESCRT-III was abundantly expressed at this time point, ESCRT-III-1 and ESCRT-III-2 were barely detectable. Conversely, at 150 min time point, when the expression of ESCRT-III-1 and ESCRT-III-2 peaked, the expression of ESCRT-III started to decline (Fig. 6b). This dynamics is consistent with the recent suggestion that ESCRT-III is the first to form a ring in the mid-cell during cell division, which serves a platform for subsequent recruitment of ESCRT-III-1 and ESCRT-III-2 [56]. We next analyzed the production of Sis-EVs in synchronized cultures at 60 (G2 phase), 90 (beginning of cell division) and 135 (advanced cell division) min after removal of acetic acid (Fig. 6c). There was a dramatic increase in EV production at the 135 min time point which coincides with active cell division (Fig. 6c, Fig. S14). These results strongly suggest that Sis-EV production is linked to the cell cycle and might be triggered by the natural, cell cycle-linked changes in the expression of ESCRT-III homologs. In particular, the active EV production appears to coincide with the expression pattern of ESCRT-III-1 and ESCRT-III-2, rather than ESCRT-III, suggesting a prime role of these proteins in EV budding. This conclusion is fully consistent with the observation that the two proteins are strongly enriched in EVs as well as with the fact that overexpression of ESCRT-III-1 and ESCRT-III-2, but not ESCRT-III, induces EV biogenesis.

**Fig. 6.**
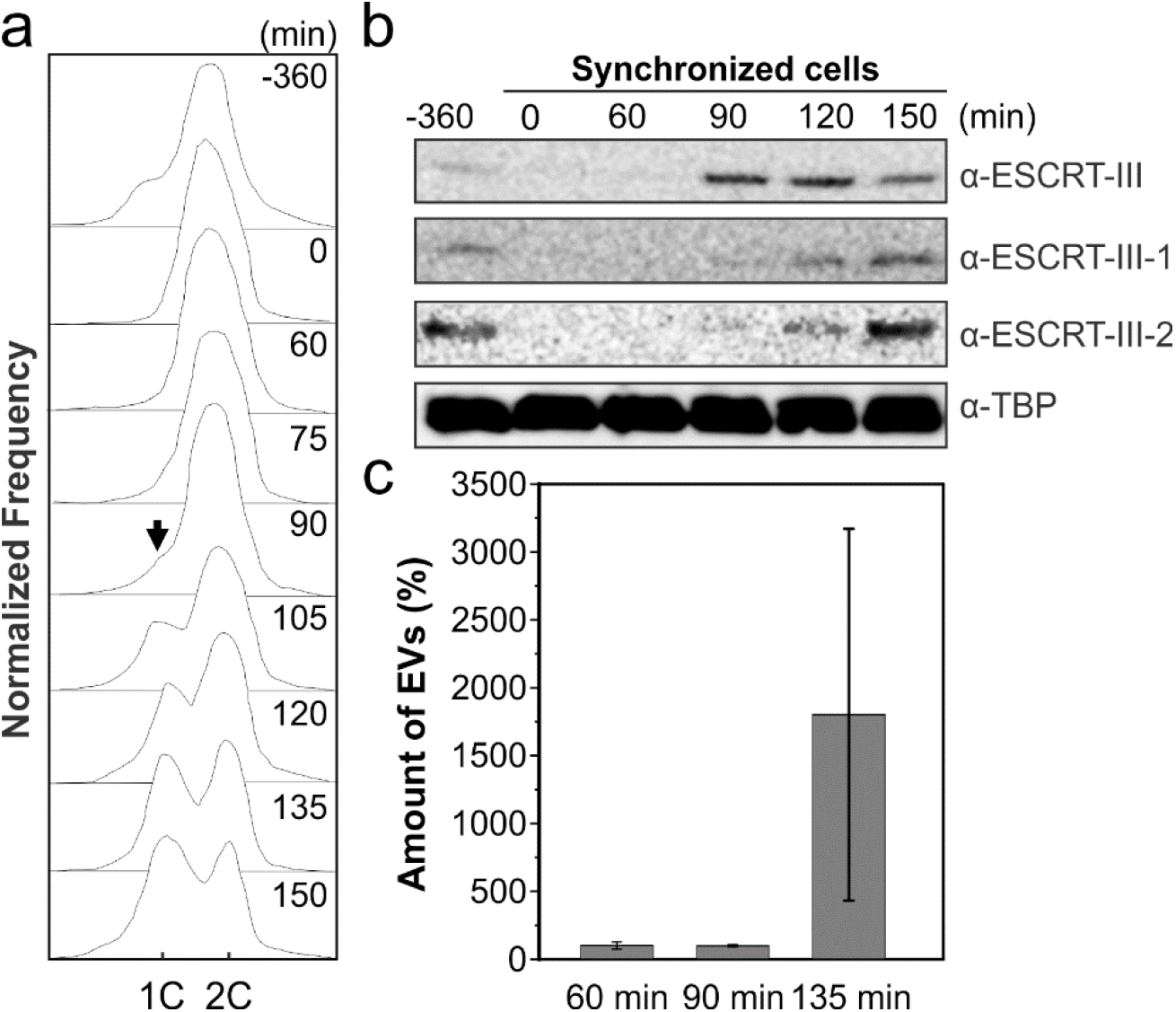
EV biogenesis is linked to cell division. **a** Flow cytometry analysis of samples taken at the indicated time points during the progression of a synchronized culture of *S. islandicus*. The positions of peaks corresponding to one chromosome copy (1C) and 2C genome contents are indicated. Black arrow indicated the reappearance of the peak corresponding to the 1C genome content, signifying cell division. **b** Western blot analysis of synchronized cells. Cells were collected at indicated time points and expression of ESCRT-III, ESCRT-III-1 and ESCRT-III-2 was analyzed using the corresponding antibodies. Tata-binding protein (TBP) was used as a loading control. **c** Flow cytometry analysis of Sis-EV production by synchronized Sis/pSeSD cells at different time points after removal of acetic acid: 60 min (prior to onset of cell division), 90 min (onset of cell division) and 135 min (active cell division). Error bars represent standard deviation from three independent experiments.

## CONCLUDING REMARKS

Here we have further characterized *Sulfolobus* EVs and showed that they carry DNA. Combined with the previous observation of DNA-containing EVs in euryarchaea (halobacteria and thermococci) [18,19,21,50,81,88,89], the finding that crenarchaeal EVs also contain DNA suggests that this property might be general across archaea. Horizontal gene transfer (HGT) is essential for the survival of microbial populations that otherwise deteriorate due to the Muller’s ratchet [90,91]. Some bacteria and archaea are naturally competent and can uptake DNA from the environment [92,93]. However, in low-density populations residing in high-temperature, acidic environments, as is the case for Sulfolobales, extracellular DNA might be neither stable nor readily available. In bacteria, conjugative plasmids, transducing bacteriophages and phage-derived gene transfer agents are considered the main drivers of the HGT. Although conjugative plasmids are known in *Sulfolobus*, their role in HGT has not been assessed [94]. By contrast, transducing viruses or dedicated gene transfer agents have not been described in Sulfolobales. Full-length genomic DNA could not be detected in the agarose gel, suggesting that only fragments of genomic DNA, which could represent byproducts of genome replication and repair, are incorporated into the Sis-EVs. Nevertheless, these DNA fragments collectively represented all genes present on the *S. islandicus* chromosome, as well as the resident plasmid. Importantly, Sis-EVs could successfully transfer the marker genes as well as the complete plasmid within the *S. islandicus* population. Furthermore, our data shows that *S. islandicus* can use EVs as carbon and nitrogen source, which is likely to be important in natural settings where nutrients are scarce. Collectively, these results indicate that EVs could play an important, yet overlooked role in gene transfer and nutrient flux in extreme environments. Indeed, we observed EVs resembling those produced by Sulfolobales directly in the environmental archaea-dominated sample, suggesting that properties of the EVs determined under laboratory conditions are biologically and environmentally relevant.

The mechanisms of EV biogenesis are poorly understood in prokaryotes [1]. Our results strongly suggest that *Sulfolobus* ESCRT machinery plays an important role in EVs formation. The finding that CdvA appears to be dispensable for this process suggests that there are mechanistic differences of the archaeal ESCRT functioning in different pathways of membrane remodeling. This would be similar to eukaryotes, where ESCRT-III complex is targeted to the membranes by different partner proteins [22]. We hypothesize that CdvA is substituted by a different targeting protein during EV budding. Notably, some archaea lack *cdvA* gene but encode ESCRT-III and Vps4 homologs [31,45-47], suggesting that ESCRT-III targeting to the membrane in these organisms, similar to eukaryotes, is achieved by an unrelated protein or proteins. Alternatively, changes in membrane curvature at the EV budding sites might promote binding of ESCRT-III paralogs, without the necessary chaperoning of CdvA. Further in vitro experiments will be necessary to test this hypothesis. Regardless, our results show that the ESCRT-dependent mechanism of EV biogenesis is conserved in both archaea and eukaryotes, and likely represents one of the ancestral functions of the ESCRT system.

## ACKNOWLEDGEMENTS

This work was supported by l’Agence Nationale de la Recherche (#ANR-17-CE15-0005-01) and Ville de Paris Emergence(s) program (project MEMREMA) grants to M.K., and European Research Council grant from the European Union’s Seventh Framework Program (FP/2007-2013)/Project EVOMOBIL-ERC Grant Agreement 340440 to P.F. J.L. was partly supported through the PRESTIGE post-doctoral program from European Union’s Seventh Framework Programme, the National Key Research and Development Program of China (No. 2020YFA0906800), and the post-doctoral fellowship from the State Key Laboratory of Microbial Technology, Shandong University. The authors would like to thank Thibault Chaze and Mariette Matondo (Proteomics Platform, Institut Pasteur) as well as Thomas Cokelaer and Marc Monot (Biomics Platform, Institut Pasteur) for help with the proteomics and sequencing analyses, respectively. We are also grateful to the Ultrastructural BioImaging (UTechS UBI) unit of Institut Pasteur for access to electron microscopes. We gratefully acknowledge the UtechS Photonic BioImaging (Imagopole), C2RT, Institut Pasteur, supported by the French National Research Agency (France BioImaging; ANR-10–INSB–04; Investments for the Future).

## CONFLICT OF INTEREST

The authors declare that they have no conflict of interest.

## Notes

### Competing Interest Statement

The authors have declared no competing interest.

## REFERENCES

1. Coelho C and Casadevall A. Answers to naysayers regarding microbial extracellular vesicles. Biochem Soc Trans. 2019; 47:1005–1012.

2. Woith E, Fuhrmann G and Melzig MF. Extracellular vesicles-connecting Kingdoms. Int J Mol Sci. 2019; 20:E5695.

3. Gill S, Catchpole R and Forterre P. Extracellular membrane vesicles in the three domains of life and beyond. FEMS Microbiol Rev. 2019; 43:273–303.

4. Brown L, Wolf JM, Prados-Rosales R and Casadevall A. Through the wall: extracellular vesicles in Gram-positive bacteria, mycobacteria and fungi. Nat Rev Microbiol. 2015; 13:620–30.

5. Schatz D and Vardi A. Extracellular vesicles -new players in cell-cell communication in aquatic environments. Curr Opin Microbiol. 2018; 43:148–154.

6. Soler N, Krupovic M, Marguet E and Forterre P. Membrane vesicles in natural environments: a major challenge in viral ecology. ISME J. 2015; 9:793–6.

7. Biller SJ, Schubotz F, Roggensack SE, Thompson AW, Summons RE and Chisholm SW. Bacterial vesicles in marine ecosystems. Science. 2014; 343:183–6.

8. Biller SJ, McDaniel LD, Breitbart M, Rogers E, Paul JH and Chisholm SW. Membrane vesicles in sea water: heterogeneous DNA content and implications for viral abundance estimates. ISME J. 2017; 11:394–404.

9. Denham J and Spencer SJ. Emerging roles of extracellular vesicles in the intercellular communication for exercise-induced adaptations. Am J Physiol Endocrinol Metab. 2020.

10. Abramowicz A and Story MD. The Long and Short of It: The Emerging Roles of Non-Coding RNA in Small Extracellular Vesicles. Cancers (Basel). 2020; 12.

11. Gho YS and Lee J. Special issue on the role of extracellular vesicles in human diseases. Exp Mol Med. 2019; 51:34.

12. Malloci M, Perdomo L, Veerasamy M, Andriantsitohaina R, Simard G and Martinez MC. Extracellular vesicles: Mechanisms in human health and disease. Antioxid Redox Signal. 2019; 30:813–856.

13. Linxweiler J and Junker K. Extracellular vesicles in urological malignancies: an update. Nat Rev Urol. 2020; 17:11–27.

14. Xu R, Rai A, Chen M, Suwakulsiri W, Greening DW and Simpson RJ. Extracellular vesicles in cancer - implications for future improvements in cancer care. Nat Rev Clin Oncol. 2018; 15:617–638.

15. Ramasubramanian L, Kumar P and Wang A. Engineering extracellular vesicles as nanotherapeutics for regenerative medicine. Biomolecules. 2019; 10.

16. Villa F, Quarto R and Tasso R. Extracellular vesicles as natural, safe and efficient drug delivery systems. Pharmaceutics. 2019; 11.

17. Gorlas A, Marguet E, Gill S, Geslin C, Guigner JM, Guyot F and Forterre P. Sulfur vesicles from Thermococcales: A possible role in sulfur detoxifying mechanisms. Biochimie. 2015; 118:356–64.

18. Erdmann S, Tschitschko B, Zhong L, Raftery MJ and Cavicchioli R. A plasmid from an Antarctic haloarchaeon uses specialized membrane vesicles to disseminate and infect plasmid-free cells. Nat Microbiol. 2017; 2:1446–1455.

19. Gaudin M, Krupovic M, Marguet E, Gauliard E, Cvirkaite-Krupovic V, Le Cam E, Oberto J and Forterre P. Extracellular membrane vesicles harbouring viral genomes. Environ Microbiol. 2014; 16:1167–75.

20. Soler N, Gaudin M, Marguet E and Forterre P. Plasmids, viruses and virus-like membrane vesicles from Thermococcales. Biochem Soc Trans. 2011; 39:36–44.

21. Choi DH, Kwon YM, Chiura HX, Yang EC, Bae SS, Kang SG, Lee JH, Yoon HS and Kim SJ. Extracellular Vesicles of the Hyperthermophilic Archaeon “Thermococcus onnurineus” NA1T. Appl Environ Microbiol. 2015; 81:4591–9.

22. Vietri M, Radulovic M and Stenmark H. The many functions of ESCRTs. Nat Rev Mol Cell Biol. 2020; 21:25–42.

23. Juan T and Furthauer M. Biogenesis and function of ESCRT-dependent extracellular vesicles. Semin Cell Dev Biol. 2018; 74:66–77.

24. van Niel G, D’Angelo G and Raposo G. Shedding light on the cell biology of extracellular vesicles. Nat Rev Mol Cell Biol. 2018; 19:213–228.

25. Christ L, Raiborg C, Wenzel EM, Campsteijn C and Stenmark H. Cellular Functions and Molecular Mechanisms of the ESCRT Membrane-Scission Machinery. Trends Biochem Sci. 2017; 42:42–56.

26. Schöneberg J, Lee IH, Iwasa JH and Hurley JH. Reverse-topology membrane scission by the ESCRT proteins. Nat Rev Mol Cell Biol. 2017; 18:5–17.

27. Hurley JH. ESCRTs are everywhere. Embo J. 2015; 34:2398–407.

28. Mierzwa BE, Chiaruttini N, Redondo-Morata L, von Filseck JM, Konig J, Larios J, Poser I, Muller-Reichert T, Scheuring S, Roux A et al. Dynamic subunit turnover in ESCRT-III assemblies is regulated by Vps4 to mediate membrane remodelling during cytokinesis. Nat Cell Biol. 2017; 19:787–798.

29. Adell MA, Vogel GF, Pakdel M, Muller M, Lindner H, Hess MW and Teis D. Coordinated binding of Vps4 to ESCRT-III drives membrane neck constriction during MVB vesicle formation. J Cell Biol. 2014; 205:33–49.

30. Samson RY, Dobro MJ, Jensen GJ and Bell SD. The structure, function and roles of the archaeal ESCRT apparatus. Subcell Biochem. 2017; 84:357–377.

31. Zaremba-Niedzwiedzka K, Caceres EF, Saw JH, Backstrom D, Juzokaite L, Vancaester E, Seitz KW, Anantharaman K, Starnawski P, Kjeldsen KU et al. Asgard archaea illuminate the origin of eukaryotic cellular complexity. Nature. 2017; 541:353–358.

32. Makarova KS, Yutin N, Bell SD and Koonin EV. Evolution of diverse cell division and vesicle formation systems in Archaea. Nat Rev Microbiol. 2010; 8:731–41.

33. Lindås AC, Karlsson EA, Lindgren MT, Ettema TJ and Bernander R. A unique cell division machinery in the Archaea. Proc Natl Acad Sci U S A. 2008; 105:18942–6.

34. Lu Z, Fu T, Li T, Liu Y, Zhang S, Li J, Dai J, Koonin EV, Li G, Chu H et al. Coevolution of Eukaryote-like Vps4 and ESCRT-III Subunits in the Asgard Archaea. mBio. 2020; 11.

35. Ng KH, Srinivas V, Srinivasan R and Balasubramanian M. The Nitrosopumilus maritimus CdvB, but not FtsZ, assembles into polymers. Archaea. 2013; 2013:104147.

36. Pelve EA, Lindas AC, Martens-Habbena W, de la Torre JR, Stahl DA and Bernander R. Cdv-based cell division and cell cycle organization in the thaumarchaeon Nitrosopumilus maritimus. Mol Microbiol. 2011; 82:555–66.

37. Samson RY, Obita T, Freund SM, Williams RL and Bell SD. A role for the ESCRT system in cell division in archaea. Science. 2008; 322:1710–3.

38. Liu J, Gao R, Li C, Ni J, Yang Z, Zhang Q, Chen H and Shen Y. Functional assignment of multiple ESCRT-III homologs in cell division and budding in Sulfolobus islandicus. Mol Microbiol. 2017; 105:540–553.

39. Yang N and Driessen AJ. Deletion of cdvB paralogous genes of Sulfolobus acidocaldarius impairs cell division. Extremophiles. 2014; 18:331–9.

40. Pulschen AA, Mutavchiev DR, Culley S, Sebastian KN, Roubinet J, Roubinet M, Risa GT, van Wolferen M, Roubinet C, Schmidt U et al. Live Imaging of a Hyperthermophilic Archaeon Reveals Distinct Roles for Two ESCRT-III Homologs in Ensuring a Robust and Symmetric Division. Curr Biol. 2020.

41. Dobro MJ, Samson RY, Yu Z, McCullough J, Ding HJ, Chong PL, Bell SD and Jensen GJ. Electron cryotomography of ESCRT assemblies and dividing Sulfolobus cells suggests that spiraling filaments are involved in membrane scission. Mol Biol Cell. 2013; 24:2319–27.

42. Samson RY, Duggin IG and Bell SD. Analysis of the Archaeal ESCRT Apparatus. Methods Mol Biol. 2019; 1998:1–11.

43. Moriscot C, Gribaldo S, Jault JM, Krupovic M, Arnaud J, Jamin M, Schoehn G, Forterre P, Weissenhorn W and Renesto P. Crenarchaeal CdvA forms double-helical filaments containing DNA and interacts with ESCRT-III-like CdvB. PLoS One. 2011; 6:e21921.

44. Samson RY, Obita T, Hodgson B, Shaw MK, Chong PL, Williams RL and Bell SD. Molecular and structural basis of ESCRT-III recruitment to membranes during archaeal cell division. Mol Cell. 2011; 41:186–96.

45. Caspi Y and Dekker C. Dividing the Archaeal Way: The Ancient Cdv Cell-Division Machinery. Front Microbiol. 2018; 9:174.

46. Abby SS, Melcher M, Kerou M, Krupovic M, Stieglmeier M, Rossel C, Pfeifer K and Schleper C. Candidatus Nitrosocaldus cavascurensis, an ammonia oxidizing, extremely thermophilic archaeon with a highly mobile genome. Front Microbiol. 2018; 9:28.

47. Nunoura T, Takaki Y, Kakuta J, Nishi S, Sugahara J, Kazama H, Chee GJ, Hattori M, Kanai A, Atomi H et al. Insights into the evolution of Archaea and eukaryotic protein modifier systems revealed by the genome of a novel archaeal group. Nucleic Acids Res. 2011; 39:3204–23.

48. Snyder JC, Samson RY, Brumfield SK, Bell SD and Young MJ. Functional interplay between a virus and the ESCRT machinery in archaea. Proc Natl Acad Sci U S A. 2013; 110:10783–7.

49. Ellen AF, Albers SV, Huibers W, Pitcher A, Hobel CF, Schwarz H, Folea M, Schouten S, Boekema EJ, Poolman B et al. Proteomic analysis of secreted membrane vesicles of archaeal Sulfolobus species reveals the presence of endosome sorting complex components. Extremophiles. 2009; 13:67–79.

50. Marguet E, Gaudin M, Gauliard E, Fourquaux I, le Blond du Plouy S, Matsui I and Forterre P. Membrane vesicles, nanopods and/or nanotubes produced by hyperthermophilic archaea of the genus Thermococcus. Biochem Soc Trans. 2013; 41:436–42.

51. Deng L, Zhu H, Chen Z, Liang YX and She Q. Unmarked gene deletion and host-vector system for the hyperthermophilic crenarchaeon Sulfolobus islandicus. Extremophiles. 2009; 13:735–46.

52. Martusewitsch E, Sensen CW and Schleper C. High spontaneous mutation rate in the hyperthermophilic archaeon Sulfolobus solfataricus is mediated by transposable elements. J Bacteriol. 2000; 182:2574–81.

53. Baquero DP, Contursi P, Piochi M, Bartolucci S, Liu Y, Cvirkaite-Krupovic V, Prangishvili D and Krupovic M. New virus isolates from Italian hydrothermal environments underscore the biogeographic pattern in archaeal virus communities. ISME J. 2020; 14:1821–1833.

54. Alkhatatbeh MJ, Enjeti AK, Baqar S, Ekinci EI, Liu D, Thorne RF and Lincz LF. Strategies for enumeration of circulating microvesicles on a conventional flow cytometer: Counting beads and scatter parameters. J Circ Biomark. 2018; 7:1849454418766966.

55. Inglis HC, Danesh A, Shah A, Lacroix J, Spinella PC and Norris PJ. Techniques to improve detection and analysis of extracellular vesicles using flow cytometry. Cytometry A. 2015; 87:1052–63.

56. Tarrason Risa G, Hurtig F, Bray S, Hafner AE, Harker-Kirschneck L, Faull P, Davis C, Papatziamou D, Mutavchiev DR, Fan C et al. The proteasome controls ESCRT-III-mediated cell division in an archaeon. Science. 2020; 369.

57. Lundgren M, Andersson A, Chen L, Nilsson P and Bernander R. Three replication origins in Sulfolobus species: synchronous initiation of chromosome replication and asynchronous termination. Proc Natl Acad Sci U S A. 2004; 101:7046–51.

58. Robertson J, McGoverin C, Vanholsbeeck F and Swift S. Optimisation of the Protocol for the LIVE/DEAD((R)) BacLight(TM) Bacterial Viability Kit for Rapid Determination of Bacterial Load. Front Microbiol. 2019; 10:801.

59. Leuko S, Legat A, Fendrihan S and Stan-Lotter H. Evaluation of the LIVE/DEAD BacLight kit for detection of extremophilic archaea and visualization of microorganisms in environmental hypersaline samples. Appl Environ Microbiol. 2004; 70:6884–6.

60. Makarova KS, Wolf YI and Koonin EV. Archaeal Clusters of Orthologous Genes (arCOGs): An update and application for analysis of shared features between Thermococcales, Methanococcales, and Methanobacteriales. Life (Basel). 2015; 5:818–40.

61. Langmead B and Salzberg SL. Fast gapped-read alignment with Bowtie 2. Nat Methods. 2012; 9:357–9.

62. Desvillechabrol D, Bouchier C, Kennedy S and Cokelaer T. Sequana coverage: detection and characterization of genomic variations using running median and mixture models. Gigascience. 2018; 7.

63. Li Y, Pan S, Zhang Y, Ren M, Feng M, Peng N, Chen L, Liang YX and She Q. Harnessing Type I and Type III CRISPR-Cas systems for genome editing. Nucleic Acids Res. 2016; 44:e34.

64. Peng W, Feng M, Feng X, Liang YX and She Q. An archaeal CRISPR type III-B system exhibiting distinctive RNA targeting features and mediating dual RNA and DNA interference. Nucleic Acids Res. 2015; 43:406–17.

65. Schmittgen TD and Livak KJ. Analyzing real-time PCR data by the comparative C(T) method. Nat Protoc. 2008; 3:1101–8.

66. Tran F and Boedicker JQ. Plasmid characteristics modulate the propensity of gene exchange in bacterial vesicles. J Bacteriol. 2019; 201:e00430–18.

67. Zhang C, Wipfler RL, Li Y, Wang Z, Hallett EN and Whitaker RJ. Cell structure changes in the hyperthermophilic crenarchaeon Sulfolobus islandicus lacking the S-layer. mBio. 2019; 10.

68. Zink IA, Pfeifer K, Wimmer E, Sleytr UB, Schuster B and Schleper C. CRISPR-mediated gene silencing reveals involvement of the archaeal S-layer in cell division and virus infection. Nat Commun. 2019; 10:4797.

69. Peng N, Han W, Li Y, Liang Y and She Q. Genetic technologies for extremely thermophilic microorganisms of Sulfolobus, the only genetically tractable genus of crenarchaea. Sci China Life Sci. 2017; 60:370–385.

70. Morales-Aparicio JC, Lara Vasquez P, Mishra S, Barran-Berdon AL, Kamat M, Basso KB, Wen ZT and Brady LJ. The Impacts of Sortase A and the 4’-Phosphopantetheinyl Transferase Homolog Sfp on Streptococcus mutans Extracellular Membrane Vesicle Biogenesis. Front Microbiol. 2020; 11:570219.

71. Rodovalho VR, da Luz BSR, Rabah H, do Carmo FLR, Folador EL, Nicolas A, Jardin J, Briard-Bion V, Blottiere H, Lapaque N et al. Extracellular Vesicles Produced by the Probiotic Propionibacterium freudenreichii CIRM-BIA 129 Mitigate Inflammation by Modulating the NF-kappaB Pathway. Front Microbiol. 2020; 11:1544.

72. Tartaglia NR, Nicolas A, Rodovalho VR, Luz B, Briard-Bion V, Krupova Z, Thierry A, Coste F, Burel A, Martin P et al. Extracellular vesicles produced by human and animal Staphylococcus aureus strains share a highly conserved core proteome. Sci Rep. 2020; 10:8467.

73. Zwarycz AS, Livingstone PG and Whitworth DE. Within-species variation in OMV cargo proteins: the Myxococcus xanthus OMV pan-proteome. Mol Omics. 2020; 16:387–397.

74. Vorontsov EA, Rensen E, Prangishvili D, Krupovic M and Chamot-Rooke J. Abundant Lysine Methylation and N-Terminal Acetylation in Sulfolobus islandicus Revealed by Bottom-Up and Top-Down Proteomics. Mol Cell Proteomics. 2016; 15:3388–3404.

75. Chu Y, Zhu Y, Chen Y, Li W, Zhang Z, Liu D, Wang T, Ma J, Deng H, Liu ZJ et al. aKMT Catalyzes Extensive Protein Lysine Methylation in the Hyperthermophilic Archaeon Sulfolobus islandicus but is Dispensable for the Growth of the Organism. Mol Cell Proteomics. 2016; 15:2908–23.

76. Ellen AF, Rohulya OV, Fusetti F, Wagner M, Albers SV and Driessen AJ. The sulfolobicin genes of Sulfolobus acidocaldarius encode novel antimicrobial proteins. J Bacteriol. 2011; 193:4380–7.

77. Prangishvili D, Holz I, Stieger E, Nickell S, Kristjansson JK and Zillig W. Sulfolobicins, specific proteinaceous toxins produced by strains of the extremely thermophilic archaeal genus Sulfolobus. J Bacteriol. 2000; 182:2985–8.

78. Schwanhausser B, Busse D, Li N, Dittmar G, Schuchhardt J, Wolf J, Chen W and Selbach M. Global quantification of mammalian gene expression control. Nature. 2011; 473:337–42.

79. Zhang Z, Guo L and Huang L. Archaeal chromatin proteins. Sci China Life Sci. 2012; 55:377–85.

80. White MF and Bell SD. Holding it together: chromatin in the Archaea. Trends Genet. 2002; 18:621–6.

81. Forterre P, Soler N, Krupovic M, Marguet E and Ackermann HW. Fake virus particles generated by fluorescence microscopy. Trends Microbiol. 2013; 21:1–5.

82. Gaudin M, Gauliard E, Schouten S, Houel-Renault L, Lenormand P, Marguet E and Forterre P. Hyperthermophilic archaea produce membrane vesicles that can transfer DNA. Environ Microbiol Rep. 2013; 5:109–16.

83. Hamm JN, Erdmann S, Eloe-Fadrosh EA, Angeloni A, Zhong L, Brownlee C, Williams TJ, Barton K, Carswell S, Smith MA et al. Unexpected host dependency of Antarctic Nanohaloarchaeota. Proc Natl Acad Sci U S A. 2019; 116:14661–14670.

84. Huber H, Hohn MJ, Rachel R, Fuchs T, Wimmer VC and Stetter KO. A new phylum of Archaea represented by a nanosized hyperthermophilic symbiont. Nature. 2002; 417:63–7.

85. Zebec Z, Manica A, Zhang J, White MF and Schleper C. CRISPR-mediated targeted mRNA degradation in the archaeon Sulfolobus solfataricus. Nucleic Acids Res. 2014; 42:5280–8.

86. Wurtzel O, Sapra R, Chen F, Zhu Y, Simmons BA and Sorek R. A single-base resolution map of an archaeal transcriptome. Genome Res. 2010; 20:133–41.

87. Zhang C, Phillips APR, Wipfler RL, Olsen GJ and Whitaker RJ. The essential genome of the crenarchaeal model Sulfolobus islandicus. Nat Commun. 2018; 9:4908.

88. Soler N and Forterre P. Vesiduction: the fourth way of HGT. Environ Microbiol. 2020; 22:2457–2460.

89. Soler N, Marguet E, Verbavatz JM and Forterre P. Virus-like vesicles and extracellular DNA produced by hyperthermophilic archaea of the order Thermococcales. Res Microbiol. 2008; 159:390–9.

90. Koonin EV. Horizontal gene transfer: essentiality and evolvability in prokaryotes, and roles in evolutionary transitions. F1000Res. 2016; 5.

91. Takeuchi N, Kaneko K and Koonin EV. Horizontal gene transfer can rescue prokaryotes from Muller’s ratchet: benefit of DNA from dead cells and population subdivision. G3 (Bethesda). 2014; 4:325–39.

92. Blokesch M. Natural competence for transformation. Curr Biol. 2016; 26:R1126–R1130.

93. Hileman TH and Santangelo TJ. Genetics Techniques for Thermococcus kodakarensis. Front Microbiol. 2012; 3:195.

94. Stedman KM, She Q, Phan H, Holz I, Singh H, Prangishvili D, Garrett R and Zillig W. pING family of conjugative plasmids from the extremely thermophilic archaeon Sulfolobus islandicus: insights into recombination and conjugation in Crenarchaeota. J Bacteriol. 2000; 182:7014–20.

